# Single-cell mutation calling and phylogenetic tree reconstruction with loss and recurrence

**DOI:** 10.1101/2022.01.28.478229

**Authors:** Jack Kuipers, Jochen Singer, Niko Beerenwinkel

## Abstract

Tumours evolve as heterogeneous populations of cells, which may be distinguished by different genomic aberrations. The resulting intra-tumour heterogeneity plays an important role in cancer patient relapse and treatment failure, so that obtaining a clear understanding of each patient’s tumour composition and evolutionary history is key for personalised therapies. Single-cell sequencing now provides the possibility to resolve tumour heterogeneity at the highest resolution of individual tumour cells, but brings with it challenges related to the particular noise profiles of the sequencing protocols as well as the uncertainty of the underlying evolutionary process. By modelling the noise processes and allowing mutations to be lost or to reoccur during tumour evolution, we present a method to jointly call mutations in each cell, reconstruct the phylogenetic relationship between cells, and determine the locations of mutational losses and recurrences. Our Bayesian approach allows us to accurately call mutations as well as to quantify our certainty in such predictions. We show the advantages of allowing mutational loss or recurrence with simulated data and present its application to tumour single-cell sequencing data.

## Introduction

The development and rapid progress in single-cell DNA sequencing [1, 2, 3] now allows the genetic profiling of individual cells. Particularly for tumours, where somatic cell evolution can lead to multiple heterogeneous cell populations and subclones [4, 5, 6], single-cell sequencing (SCS) illuminates the underlying complexity or intra-tumour heterogeneity (ITH) [7]. Measuring and understanding ITH is central for precision medicine, given its strong links to tumour relapse and treatment failure [8, 9, 10, 11].

The power and resolution of SCS come with the cost of elevated error rates, due to the small amount of DNA present in each individual cell [1, 2, 3]. For whole-exome sequencing (WES), a common amplification protocol is multple-displacement amplifiction (MDA) [12], which is efficient in creating enough DNA material for later sequencing, but suffers from uneven coverage and a high (10%–20%) rate of allelic dropout whereby one allele is locally not amplified at all and cannot be later detected in the sequencing data.

If the SCS data is dichotomised into mutational presence or absence per cell, a suite of phylogenetic methods have been developed [13, 14] to handle the high false negative rates (due to allelic dropout) particular to SCS and accurately reconstruct the evolutionary history of tumours from the genetic profiles of individual cells.

In order to obtain dichotomised data, the mutations need to be called per cell based on the raw sequencing output. Bulk callers adapted to the noise profiles of the mixed signals of many cells amplified with a different protocol are suboptimal, which has led to the development of specialised callers for single-cell data [15, 16] accounting for the noise profiles of single-cell amplification and sequencing protocols. They often share information across cells [15] or locally across the genome [16, 17] to improve performance. Combining single-cell-specific read count modelling with single-cell phylogenetic modelling, we previously developed SCIΦ [18] to jointly call mutations and learn the lineage relationships between cells. As a Bayesian approach, the full posterior certainty in the mutation calls can be assessed. The tree structure allows information to be shared more effectively across cells, particularly in correcting for allelic dropout [18] leading to improved performance as compared to combining information across cells without using the phylogeny [15].

The underlying tree model for SCIΦ contained the simplifying *infinite sites assumption* which restricts mutations to only occur once in the phylogeny and to persist after occurrence, though the model did allow for homozygous mutations. Binarised single-cell data has allowed us to test such assumptions and find that it may be often violated in real tumour samples [19]. More complex phylogenetic models mitigating or avoiding entirely the infinite sites assumption have also been developed [20, 21, 22, 23, 24], though there is an apparent trade-off in model complexity between too simple models that cannot capture all relevant aspects of the evolutionary process and too complex models that are prone to overfitting or computationally too expensive to be learned efficiently from data. The existing models rely on processed data, where the mutations have already been called. Here, we therefore bring the advances of allowing mutational recurrence and loss in tree modelling to improve mutation calling from raw singlecell sequencing data. We present a novel approach to relaxing the infinite sites assumption, building on SCIΦ [18] while staying within the same computational complexity class. The new method, called SCIΦN^‡^, allows us to jointly call mutations and the phylogenetic relationship between cells under loss and recurrence, while quantifying the uncertainty in our results.

## Results

### Model overview

During tumour evolution, mutations may be accumulated by cells, but regions of the genome may also undergo copy number changes, in particular the loss of one allele (loss of heterozygosity, LOH). In our model, SCIΦN, we consider originally diploid regions of the genome which may experience somatic point mutations, monoploid regions which have already lost one allele, as well as the loss of one allele, including its mutations, during tumour evolution (Figure 1).

**Figure 1:**
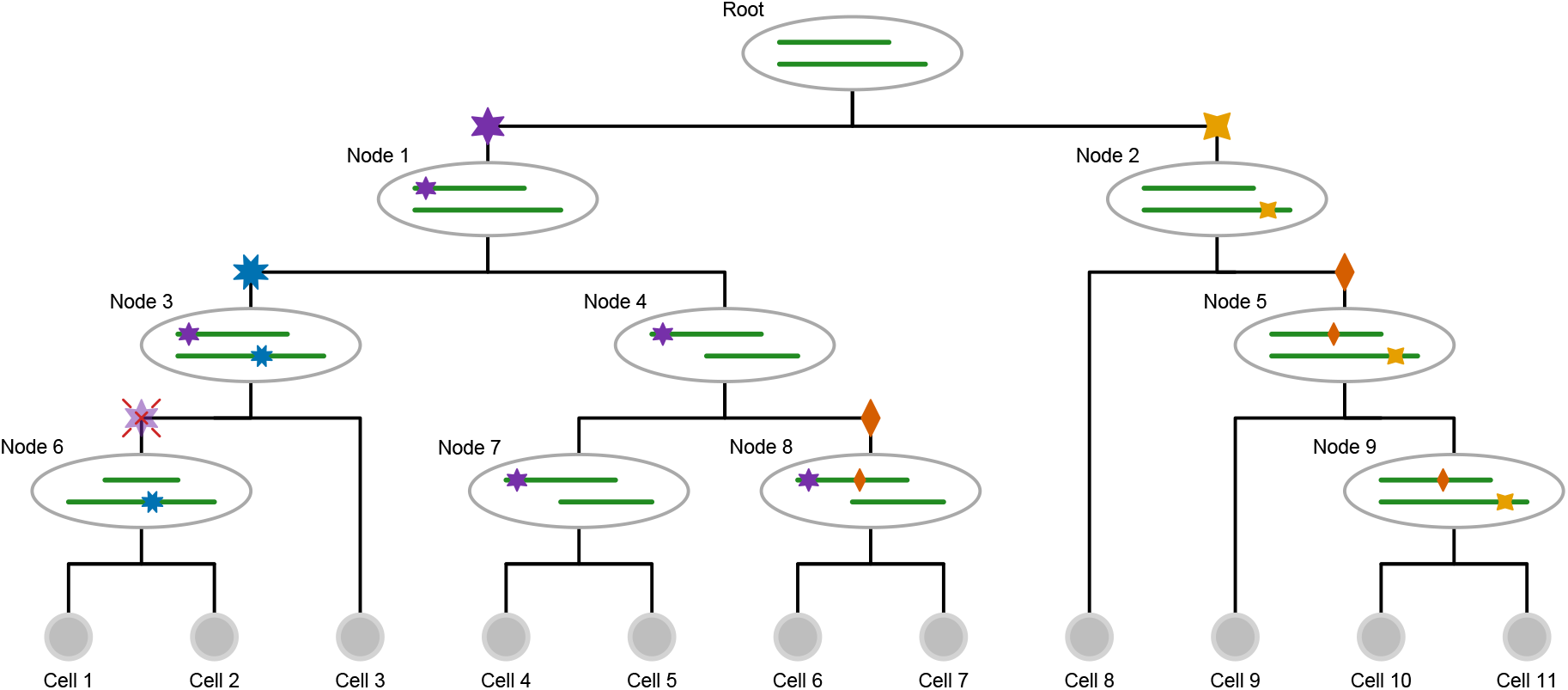
Genomic events modelled in the cell lineage tree. Starting from the tumour founder cell at the root which has a deletion in one genomic region, different clones evolve along the different lineages. On the right hand branch at node 2, the yellow (four-pointed star) mutation occurs as a *hemizygous* mutation in the remaining allele of the deleted region, later joined at node 5 by the *heterozygous* red (two-pointed star) mutation. The mutation reoccurs independently at node 8. There are two loss of heterozygosity events: In the left branch, the heterozygous purple (six-pointed star) mutation at node 1 becomes hemizygous in its right subtree at node 4 when the non-mutated *wild type* allele region is lost. At node 6 in its left subtree instead, the allele carrying the purple (six-pointed star) mutation is lost so that the mutation status returns to wild type.

At each genomic position, a cell may then be *wild type*, or have a *heterozygous* or *hemizygous* mutation. Along with mutational losses, mutations are also allowed to reoccur independently in the phylogeny. With this space of underlying aberration events, we develop the probabilistic tree model for single-cell read and variant counts, and employ MCMC to perform Bayesian inference of mutation calls.

A cell lineage tree *T* is a binary tree with labelled leaves corresponding to the single cells. Along the branches of the tree, mutational events may occur. For each genomic locus *i*, we record as element *τ_i_* of the vector *τ* the branch where the mutation affecting that locus occurs. For example, the blue 8-pointed star mutation in Figure 1 occurs in the branch above node 3 and is present in all descendant cells (cells 1, 2 and 3). The knowledge of the tree *T* and the placement of the mutations within that tree *τ*, provides us with the underlying genotypes of each cell *j*. For each genomic locus *i* and cell *j* with *v_ij_* variant reads and coverage *c_ij_*, we summarize the data as *D_ij_* = (*v_ij_, c_ij_*). This allows us to define the likelihood of the data

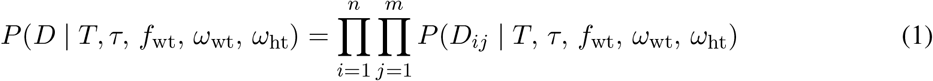

where *n* is the number of genomic loci, *m* the number of cells and we assume independence of the noise per cell and mutation. The parameters *f*_wt_, *ω*_wt_, and *ω*_ht_ are related to the noise modelling of the amplification and sequencing protocols which we expound in the Methods section. For notational convenience, we drop their explicit dependence in the following.

For Bayesian inference of the tree topology and mutation calls, we first marginalise over the unknown placement of the mutations (for which we use a uniform prior over the tree branches)

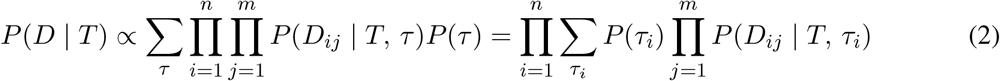

By rearranging the terms to treat each mutation separately, we reduce the complexity of the computation from the naïve *O*(*nm*^*n*+1^) on the left of the equality to *O*(*nm*^2^) on the right. As shown in the Methods section, this can be further reduced to *O*(*nm*) via tree traversals.

To compute the contribution to the likelihood from each mutation, we treat five mutation cases:

ht: A heterozygous mutation occurs in a diploid region
hm: A hemizygous mutation occurs in a monoploid region which has previously lost one allele
wl: A wild type allele is lost after a heterozygous mutation occurred
ml: A mutated allele is lost after a heterozygous mutation occurred
pm: A heterozygous mutation occurs twice in the tree in parallel branches

We consider that each possible mutation type has a fixed prior probability, namely *v* for a mutation occurring in a genomic region with only one allele, λ_wl_ and λ_ml_ for losses of heterozygosity, and *κ* for a parallel heterozygous mutation. We may then express the likelihood contributions as a mixture of the possibilities

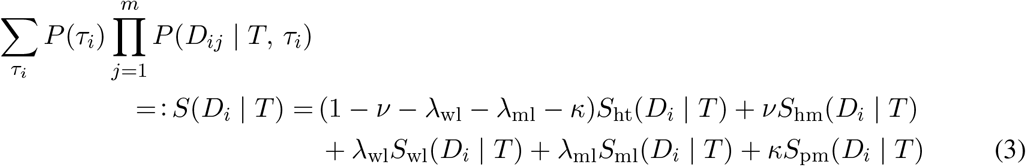

We detail in the Methods section how to compute the individual terms *S* in this mixture in time *O*(*m*) using tree traversals and tracking partial sums. The overall time complexity of computing the tree marginalised likelihood is *O*(*mn*), the same complexity class as in SCIΦ.

Artefacts in sequencing data may mimic the effects of violations of the infinite sites assumption [19], while their effect on the likelihood would scale with the number of mutations. To compensate for such effects, we introduce a regularising prior, compounded for each lost or parallel mutation, with an exponential form as

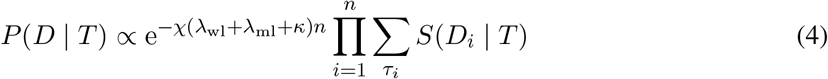

where *χ* = 0 would correspond to no regularisation while the limit *χ* → ∞ would enforce the infinite sites assumption and allow no lost or parallel mutations. A fixed prior, which does not scale with the number of mutations, would roughly correspond to 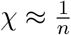.

### Benchmarking on simulated data

To explore the performance in calling mutations we simulated data with various violations of the infinite sites assumption (Figure 2). First, we added increasing amounts of losses with no parallel mutations (Figure 2a and Supplementary Figure S1), and we see similar F1 performance to SCIΦ, with better performance at higher levels of losses, but a slight decrease in performance when there are no losses. The more complicated model of SCIΦN can explain some randomly correlated drop-out events as mutational losses, which is ruled out by the model of SCIΦ leading to a slight relative loss of recall, but better precision since actual loss events can now be properly identified by SCIΦN instead of being misclassified by SCIΦ. Loss of wild type alleles results in hemizygous mutations, which are harder to misclassify as unmutated, leading to the general increase in recall at higher rates of loss (Supplementary Figure S1a).

**Figure 2:**
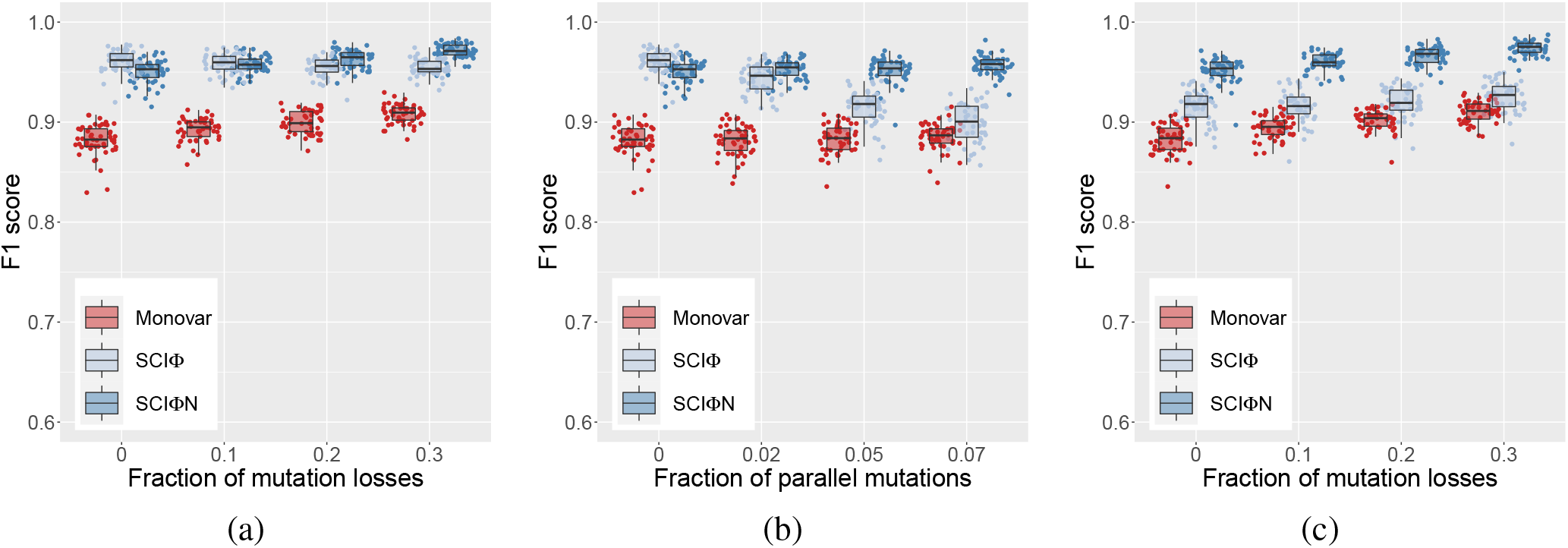
Effect of infinite sites violations on single cell mutation calling. (a) Losses, with no parallel mutations. (b) Parallel mutations, with no losses. (c) Losses with 5% parallel mutations.

With parallel mutations (Figure 2b and Supplementary Figure S2) there is a clear degradation in the precision of SCIΦ, while SCIΦN has near perfect precision because such events are included in the modelling. There is a slight cost of the more complex model of SCIΦN in terms of the recall, but the overall F1 score show the advantage of the SCIΦN model even at low rates of parallel mutations.

When both losses and parallel mutations are present (Figure 2c and Supplementary Figure S3) these effects combine to amplify the improvement of SCIΦN over the simpler infinite sites model of SCIΦ. In all cases (Figure 2) SCIΦN performs more strongly than Monovar [15] since Monovar does not use the phylogenetic relationship between cells to help improve mutation calling.

### Mutation calling and phylogenetic reconstruction from tumour data

First we applied SCIΦN to a whole-exome sequencing dataset of 16 single cells from a breast cancer [25]. For the somatic mutations previously identified by SCIΦ, we examine the effect of the regularisation of losses and parallel mutation (Figure 3). This can be seen more clearly when we separate out the contributions to the probability of mutation presence from the different mutation types considered in our modelling (Figure 4 and Supplementary Figure S4). While the majority of mutations are shared in all cells (with the possible exception of cell h1 where mutation calling is more uncertain) we observe significant amounts of loss with no penalisation (**χ** = 0, Figure 3a, Supplementary Figure S4 top row). With increasing penalisation only losses and parallel mutations with stronger evidence in the sequencing data are retained until none are allowed under the infinite sites assumption (*χ* = ∞, Figure 3e, Supplementary Figure S4 bottom row).

**Figure 3:**
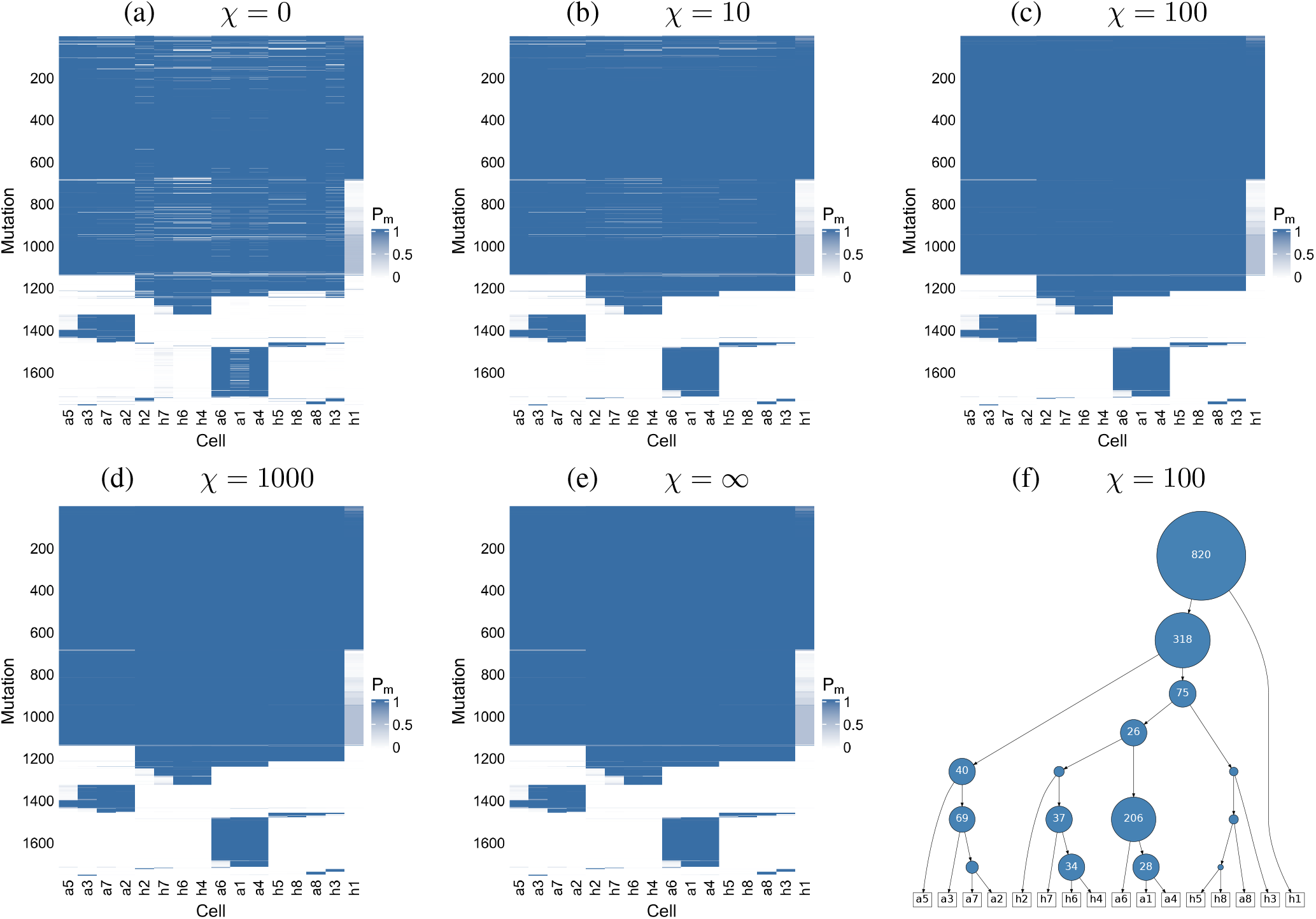
Mutation calling on 16 breast cancer cells. (a – e) the probability of mutation presence, *P_m_* in the single cells for different values of the regularisation *χ* on parallel mutations and mutation losses. The ordering of the mutations and cells is determined by the tree (f) learned under moderate penalisation (*χ* = 100).

**Figure 4:**
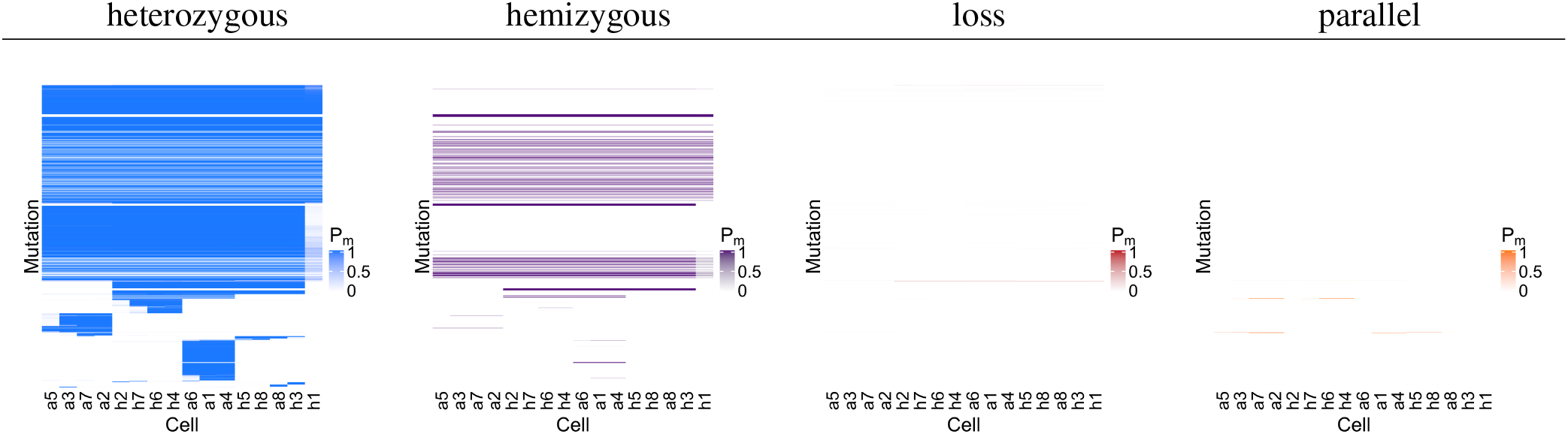
Mutation subtype calling on 16 breast cancer cells. The probability of mutation presence, *P_m_* displayed in Figure 3c with penalisation *χ* = 100, broken down into components coming from heterozygous mutations, hemizygous mutations, loss of the wild type allele or loss of the mutation elsewhere in the tree, and from parallel mutations.

When we compare the mutation calls to Monovar [15], SCIΦN finds many more mutations (Supplementary Figure S5) particularly since it can correct for allelic dropout by sharing information across cells through the inferred phylogeny, in line with the simulation results. SCIΦN and SCIΦ are highly consistent across the cells. Since SCIΦ cannot model mutational loss it must call as present mutations which are supported by other cells in a subtree even without variant reads. Whereas, for neighbouring cells without variant reads, SCIΦN can identify a shared mutational loss allowing it to call a few fewer mutations than SCIΦ. In terms of runtime, Monovar is notably faster than SCIΦN since it does not use or infer a phylogeny, taking around 1hr for this dataset compared to about 10hrs for SCIΦN (and about 6hrs for SCIΦ).

As a second dataset we considered panel-based sequencing of 255 single cells on positions detected in bulk whole-exome sequencing for a patient with acute myeloid leukemia [26]. This dataset involves high-throughput sequencing which may be more error and doublet (inadvertent sequencing of two cells together) prone and, with a large number of cells relative to the number of mutations profiled, is challenging for cell lineage reconstruction. With no penalisation (Figure 5a), we observe lots of violations of the infinite sites assumption to explain the data, which are smoothed out with moderate penalisation (Figure 5b). Under the infinite sites assumption (Figure 5c), mutations are missing which could otherwise be explained as loss or parallel mutations with more moderate penalisation (Supplementary Figure S6). The overall mutation probabilities are still highly similar across the different penalisations (Figure 5), but the assignment of their constitute parts to different mutation signatures varies significantly (Supplementary Figure S6). With no penalisation the data can be explained under the loss and parallel mutation models, while under the infinite sites model everything is explained as heterozygous mutations.

**Figure 5:**
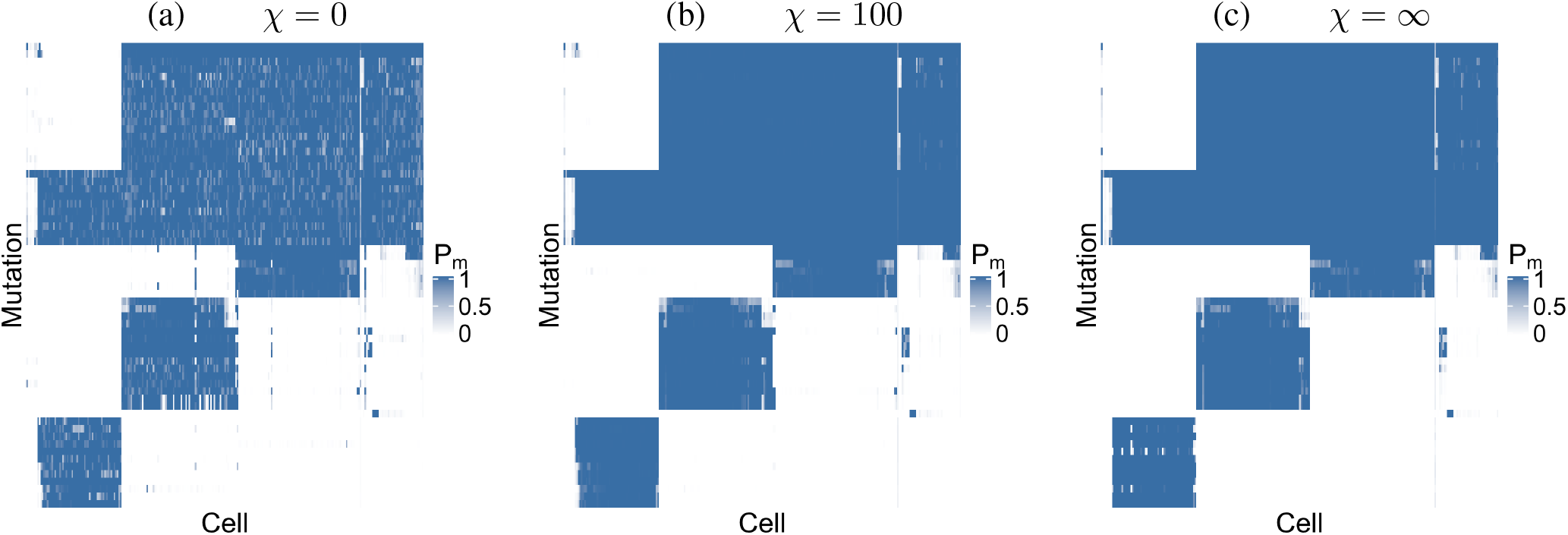
Mutation calling on 255 acute myeloid leukemia cells. (a – c) the probability *P_m_* of mutation presence in the cells as the penalisation **χ** is varied. The ordering of rows and columns is fixed to match panel (b).

In terms of runtime, Monovar is again much faster, taking around 45mins for this second dataset compared to about 30hrs for SCIΦN (and about 20hrs for SCIΦ), but does not benefit from sharing information across cells that we leverage with the phylogenetic modelling of SCIΦN.

## Conclusions

We developed SCIΦN, a tree inference method for cell lineage building and mutation calling from SCS data which allows for mutational losses and recurrent mutations, and which therefore better models the complex evolution of tumours. Compared to the previous, simpler model of SCIΦ which assumes the infinite sites assumption, the new method developed here offers superior mutation calling. Also, despite the strength of SCIΦN in modelling tumour evolution more realistically, we managed to constrain its computational complexity to the same class as the infinite sites model through tracking partial likelihood terms through judicious tree traversals.

SCIΦN considers the full read and variant counts for each cell at each genomic position to better distinguish mutations from sequencing and amplification noise. In addition, the tree building allows us to effectively share information across cells, especially to correct for allelic dropout, and improve mutation calling. In relaxing the infinite sites assumption, we only allowed certain types of violations including mutational losses and recurrent parallel mutations. Allelic dropout is relatively common in SCS data, so that violations like mutational losses in individual cells that could be easily explained by allelic dropout instead, were excluded. Likewise we also excluded violations that would recreate genotypes allowed under the infinite sites assumption. For example a pair of parallel mutations in child branches generates the same genotypes as a single mutation in the parent branch. Our relaxation therefore only considers violations which should have an additional signal in the data beyond the infinite sites base model and typical sequencing noise. The relaxation is correspondingly more conservative than transition-based classical phylogenetic models adapted for single-cell sequencing [20, 24].

Even with our stricter model, extra noise sources in the data can mimic infinite sites violations and create spurious signals [19]. For real data analyses, we include additional penalisation to reduce fitting such patterns and obtaining overly complex evolutionary histories. Though we might expect to see some violations in the infinite sites assumption during tumour evolution, we may not expect large numbers suggesting that some penalisation is required. Future work which models all noise intrinsic in the generation of SCS data will be needed to remove such penalisation. This will be particularly important for high-throughput sequencing with potentially higher noise, including doublet samples which combine genotypes from different phylogenetic branches, and relatively more cells than mutations.

For large scale datasets, the speed of MCMC schemes can become an issue. An interesting avenue has been to recode the maximum likelihood point estimate corresponding to SCIΦ as ILP constraints, which can offer significant speed-ups [27], as can hill-climbing in the search space [28]. Branch and bound algorithms have also been shown to offer a substantial speed-up for the binarised phylogeny problem [29]. Interfacing these ideas may provide pathways to speed up Bayesian inference to account for model uncertainty, as well as for the more complex model developed here.

## Methods

### Variant read model

The MDA process [12] is akin to a Pólya urn where successfully amplified genomic fragments are returned to the pool to potentially be amplified in the next round. For the distribution of variant reads at a given coverage *c*, we therefore employ a Beta-binomial distribution with density

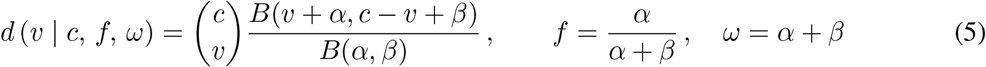

where *B* is the beta function, and we parameterise in terms of the expected frequency *f* of the variant and overdispersion *ω*. For heterozygous mutations, we expect a frequency 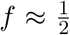. For wild type positions, we only expect sequencing or MDA errors, which can also be modelled with a Beta-binomial distribution with a low frequency *f*_wt_. When the underlying state is wild type, the likelihood is

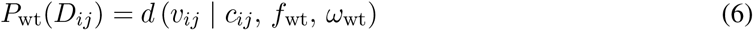

Similarly, when the underlying state is hemizygous so that only a mutated allele remains, we have

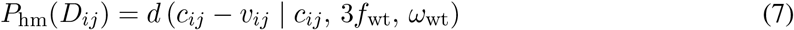

where sequencing errors may lead to any of the other 3 bases giving the additional factor here (which was not considered in [18]).

Finally, when the underlying state is heterozygous, we explicitly include an allelic dropout parameter *μ* and have the following mixture

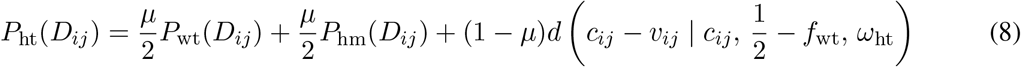

where the first two terms are the loss of the variant and reference allele, respectively, and the third when both alleles are amplified. In that case, we adjust the expected frequency slightly to account for errors resulting in the other bases (this is a corrected version compared to [18]).

### A heterozygous mutation

We compute a mixture over the different types of mutation placed upon each branch of the tree, so we start by considering, for a particular locus *i*, placing a heterozygous mutation everywhere in the tree *T* (Figure 6a) and wish to compute

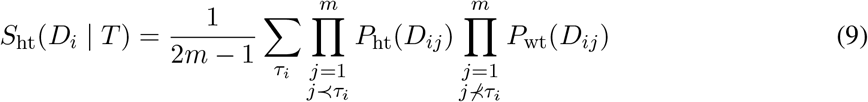

where 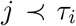 means that cell *j* is below the attachment point of the mutation and so should exhibit a heterozygous mutation at locus *i*, while it should be wild type otherwise. To proceed, we factor out the contribution where every cell is wild type

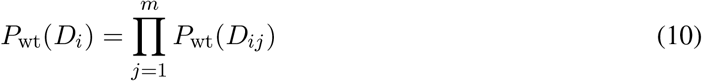

and define

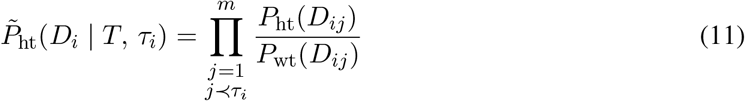

as the (relative) likelihood of the data when a heterozygous mutation at locus *i* is placed at position *τ_i_* in the tree *T*. For simplicity, we will number the branches above the leaves with the number of the cell below, from 1 to *m* and label the inner branches from (*m* +1) to (2*m* – 1), see Figure 6a. For the leaves in the tree, the computation just involves the likelihood ratio of heterozygous mutations and wild type for each cell

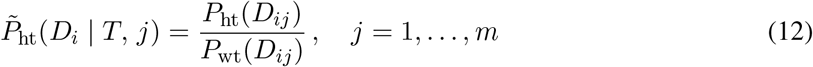

**Figure 6:**
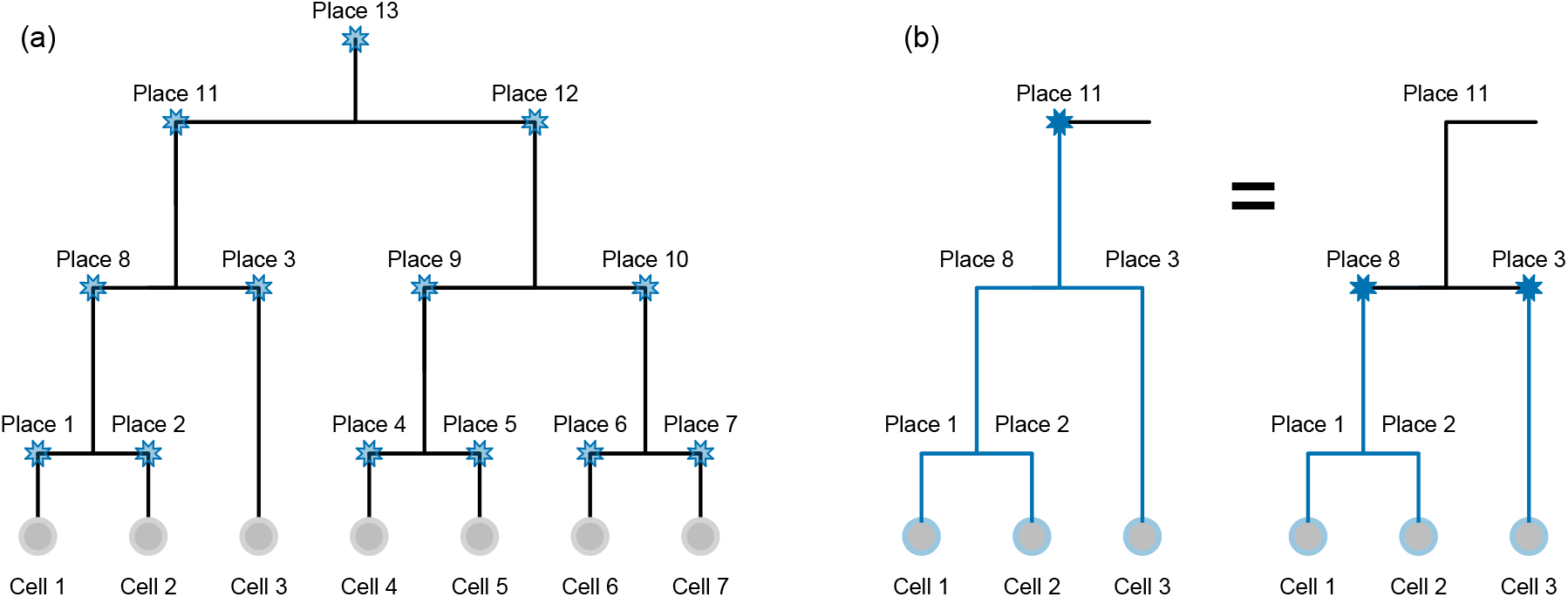
Mutations in the cell lineage tree. (a) When a mutation occurs in tumour evolution it affects all descendant cells so the possible placements of the mutation in the tree are at all inner branches. (b) A mutation occurring in the tree (for example at the place labelled 11) has equivalent genotypes at the cells as two mutations occurring along the child branches (3 and 8 in this example). The likelihood contribution of a mutation at a particular branch is therefore computed from the contributions of its children using a tree traversal.

For the inner nodes, we can compute the probabilities using a tree traversal

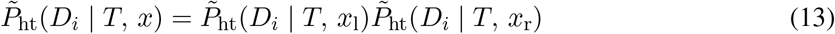

where we denote the two children of *x* in the tree *T* as *x*_l_ and *x_r_*. This relationship is illustrated in Figure 6b where having a mutation placed on the branch labelled 11 means the mutation is inherited down the tree into cells 1–3. Placing the mutation at each of the child branches (labelled 8 and 3 in Figure 6b) also ensures that cells 1–3 inherit the mutation. The likelihood contribution from having the mutation in cells 1 and 2 was computed when placing the mutation in the branch labelled 8, while the contribution from cell 3 was computed when placing the mutation in the branch labelled 3. By simply combining these two child contributions according to Equation (13) we obtain the likelihood contribution when placing the mutation at the parent branch. The sum

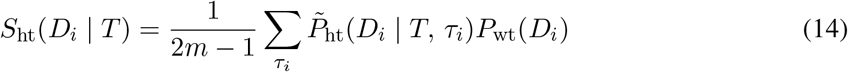

can then be computed in time *O*(*m*).

### A hemizygous mutation

If a mutation occurs in a region with only one allelic copy, we have the same recursion

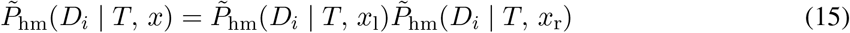

for the inner nodes and the following starting values

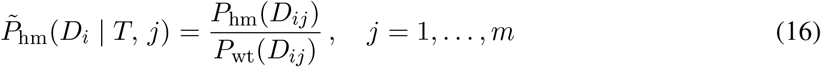

for the leaves. For the sum, however, we exclude hemizygous mutations occurring in a single cell. The rationale is that we cannot distinguish between a hemizygous mutation in a single cell from the dropout of the wild type allele in the amplification process for that cell. Since drop-out occurs relatively frequently in single-cell sequencing, we assume this as the simpler explanation of the data, and only consider hemizygous mutations when corroborated by at least two cells. We therefore define the sum as

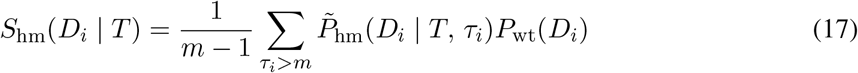

which we can again compute in time *O*(*m*).

### Loss of the wild type allele

Moving beyond the infinite sites assumption, we now consider cases where alleles are lost after mutations occur. The simplest case is when, after a heterozygous mutation, the wild type allele is lost so that the mutation becomes hemizygous. This means that in the tree, we have a heterozygous branch with a hemizygous subtree. To account for this we need to track the location of the original heterozygous mutation along with the location of the loss (Figure 7a), leading to a quadratic (in *m*) number of possibilities. However, for each location of the original heterozygous mutation, we can compute all the partial sums

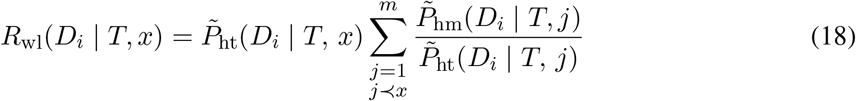

with ‘wl’ standing for *wild type loss*. For the computation we use the recursion

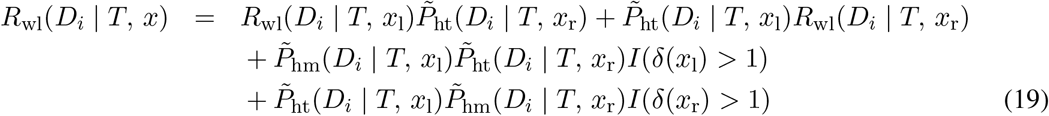

with a single tree traversal. The first line represents the case that the loss does not occur at one of the children of *x* making one subtree heterozygous and the loss occurring somewhere in the other subtree (Figure 7b,c). The next two lines represent the cases that the loss occurs directly at either of the two children of *x* so that a hemizygous and heterozygous subtree meet at *x* itself. In the recursion, we again do not allow a hemizygous mutation above a leaf, so the loss of the wild type allele must affect at least two cells, indicated in the equation by the indicator function *I* on the number of descendants *δ*(*x*). The recursion starts at 0 for each leaf:

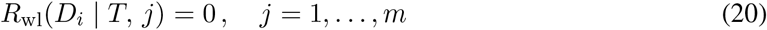

**Figure 7:**
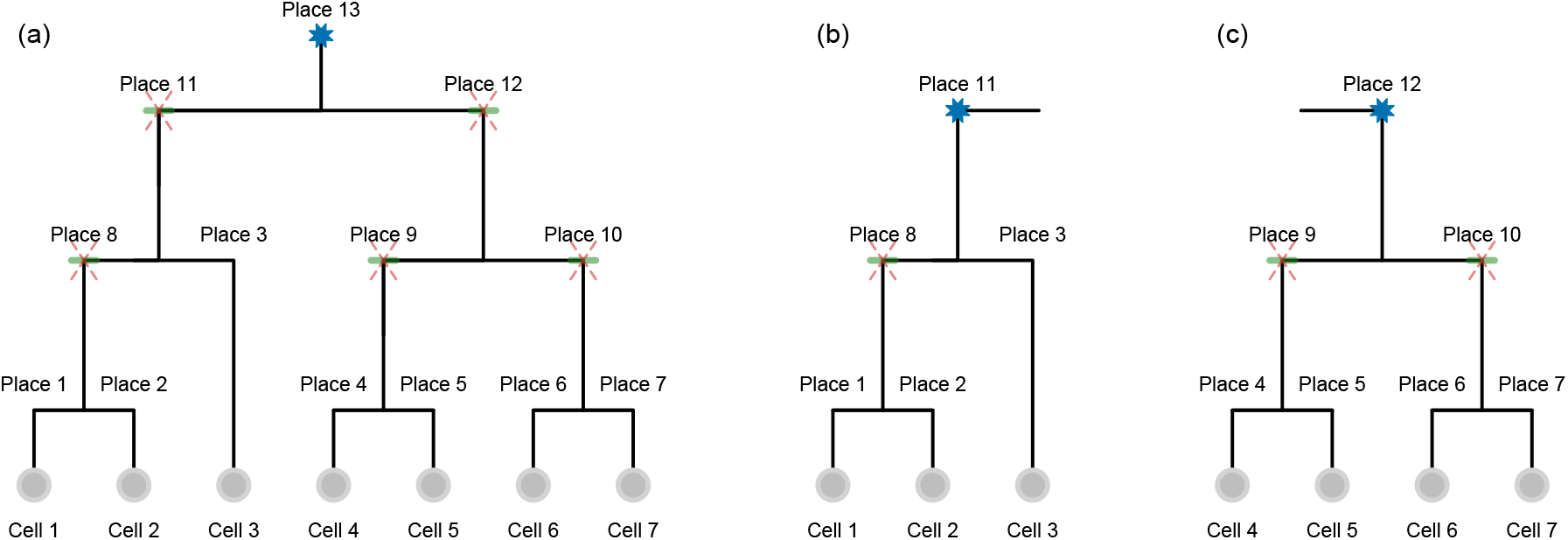
Loss in the cell lineage tree. (a) For a given placement of the original mutation (here at the place labelled 13), we wish to count all highlighted possible placements of the loss of the wild type allele (which are above at least two cells). Along with placing the loss directly at a child branch (11 and 12 in this example) the possible placements lower down in the tree are covered by the contributions of the two child subtrees (b) and (c) which have already been computed in the tree traversal.

When we sum the different possibilities

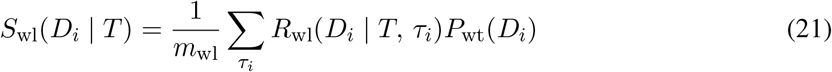

we normalise by *m*_wl_ the total number of permissible placements of the original mutation and the loss which can be counted with the same recursion as above by keeping track of the non-zero terms. Because of the recursion, even though there are a quadratic number of terms in the sum, we can evaluate in time *O*(*m*).

### Loss of the mutated allele

Alternatively we may lose the mutated allele so that in the tree there is a heterozygous region with a wild type subtree. Again we track the location of the original mutation

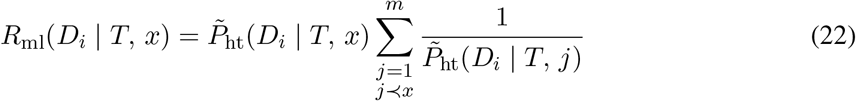

with ‘ml’ standing for *mutation loss*. This is computed using the recursion

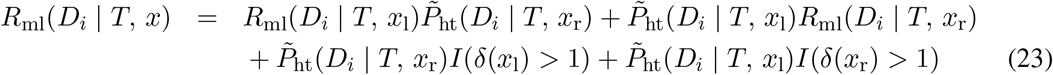

where the wild type terms just give a factor of 1 because scores are relative to the wild type case. The definition above allows for the loss of the mutated allele directly after it appears in the tree: when the mutation occurs at node *x* it is lost at node *x*_l_ or *x*_r_. A more parsimonious explanation, which leads to the same observed data, is that the mutation was gained directly at *x*_r_ or *x*_l_ instead. We therefore exclude such scenarios and define

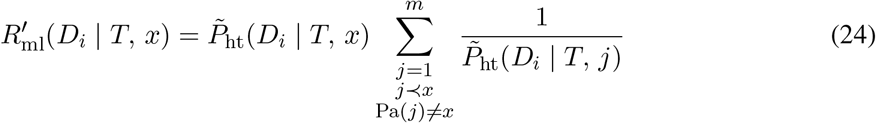

which can be computed as

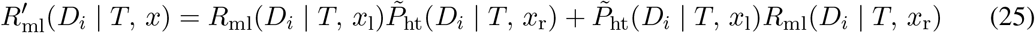

We sum the possibilities as

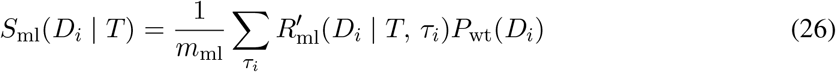

where *m*_ml_ is the total number of permissible placements of the original mutation and the loss.

### Parallel mutations

For parallel mutations we track all cases where a mutation occurs in two distinct subtrees below any given node

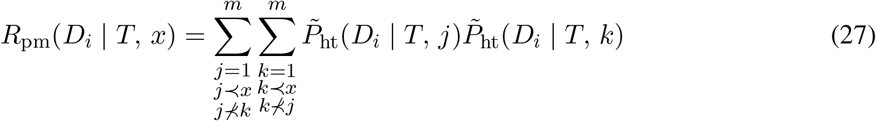

To evaluate these sums, we first compute the partial sum that a heterozygous mutation occurs somewhere in the subtree below a node *j*

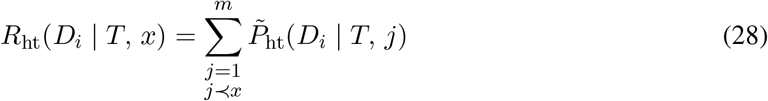

which is again computed using a tree traversal

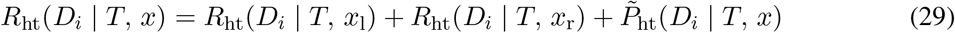

since either the mutation is lower down one branch, or occurs at *x* itself.

The parallel mutation term is

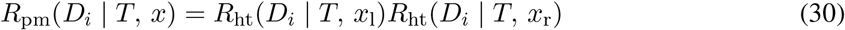

but this again includes terms with unnecessary complexity. For example the case where the mutation occurs at both *x*_l_ and *x*_r_ can be more parsimoniously explained as the mutation occurring once at *x* instead. Another case is when the mutation occurs at one of the children of *x* as well as one of the grandchildren on the other branch. This could be explained as a mutation at *x* and then a loss at the non-mutated grandchild. We therefore exclude these possibilities and define

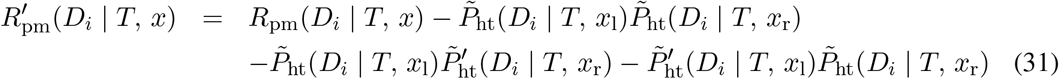

with

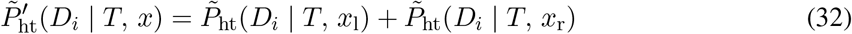

We sum the possible placements of parallel mutations as

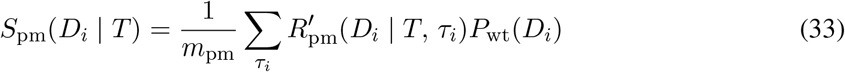

where *m*_pm_ is the number of permissible placements of both mutations (corresponding to those with a non-zero score in *R*′). The time complexity is again *O*(*m*).

### Tree scoring complexity

The overall tree score can therefore be computed in time *O*(*mn*) since we employed tree traversals to compute the terms and partial sums needed for its computation. This is akin to the peeling algorithm of Felsenstein [30] used to track partial likelihoods and marginalise the inner node states in leaf-labelled trees. The difference here is in the kind of biological effects we permit in our tree, and that our restrictions span generations leading to more complicated tree recursions. The restrictions we impose are such that if there is a simpler model which generates the exact same cell genotypes as the more complex one, we rule out the more complex case. For example if two parallel mutations can be replaced by a single mutation affecting the same cells in the tree we do not allow the parallel mutation case. Likewise if the loss of mutation can be replaced by allelic dropout, we exclude the loss from the modelling.

### Posterior mutation probabilities

With non-informative priors on the parameters and *T* we obtain *P*(*T*, *f*_wt_, *ω*_wt_, *ω*_ht_ | *D*) ∝ *P*(*D* |*T*, *f*_wt_, *ω*_wt_, *ω*_ht_). To obtain a sample from the posterior space, we employ MCMC where we may swap leaf labels or prune and reattach a subtree or perform a Gaussian random walk for the continuous parameters, as detailed in [18].

From the sample of trees and parameters, we reverse the marginalisation to obtain the posterior probability of each mutation occurring in each single cell. We average over the full sample of trees and parameters to obtain the posterior mutation probabilities.

From each sampled tree and parameter combination in the average, we first compute the probability of each mutation type. For heterozygous mutations, we know the relative probability of the mutation occurring at each node by normalising 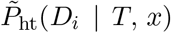 by their sum. Propagating these probabilities down from the root to the leaves provides the conditional probability of mutational presence in each cell given that it is a heterozygous mutation. For hemizygous mutations and the loss of the wild type allele we compute the probabilities analogously.

For the loss of the mutated allele, however, all we know are the relative probabilities that the mutation occurred at node *x*, but we have not recorded where the loss occurred. To remedy this, we also compute *Q*_ml_(*D_i_* |*T, x*), the probability that the mutational loss itself occurred at *x*, and we sum all possibilities that the original mutation occurred at an ancestor of *x*. To perform the tree traversal from the root down, we define *x*_p_ as the parent of *x* and *x*_s_ as its sibling, so that we can use the recursion

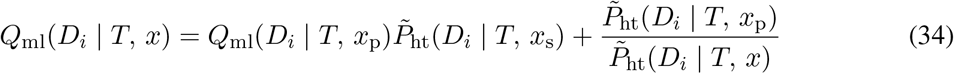

where the first term represents shifting the loss from a parent to the child and setting the other sibling to be heterozygous, while the second term is the additional case where the mutation occurs at the parent and is directly lost at one child. These need to be computed for the recursion to function, but excluded from the final set of possibilities as

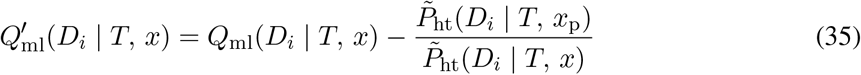

The other case is when a cell is below a parallel mutation. To compute this we track the sum of possibly placements of a parallel mutation on alternative subtrees on the path to the root

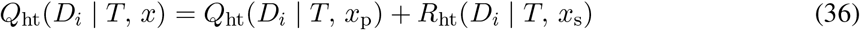

We must also remove the restricted cases when the sibling of the parent, or a child of the sibling are mutated

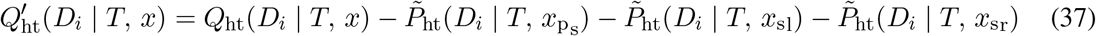

Placing the other mutation at *x* then gives the contributions

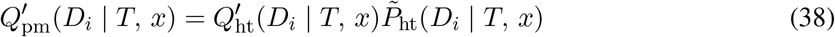

from which we again normalise and propagate down the tree to obtain the conditional probabilities of a mutation being present in each cell.

### Simulation settings

The simulated datasets were generated similar to the process described in [18]. In a first step, we generated a random binary genealogical cell linage tree with 25 cells and assigned 100 mutations to the inner nodes. The mutations of node *x* were then propagated to the leaves in the subtree rooted at x. In addition, with probability 0.2 either the mutation or the wild type allele was lost, simulating drop out events.

The mutations were then mapped to a 1 million base pair (bp) long random reference, with a nucleotide distribution following a Pòlya urn model as detailed in [18]. In addition, we also simulated sequencing errors with a frequency of 10^−3^ and PCR amplification errors with a frequency of 5 × 10^−7^. In contrast to [18], we also simulated the loss of one allele of a mutation and the appearance of the same mutation twice independently in two distinct subtrees. With probability *λ* a mutation in the subtree rooted at node *x* becomes wildtype or homozygous alternative in the subtree rooted at node *y*. Here, the choice of node *y* is uniform in the subtree of *x*. With probability *κ* two nodes in two distinct subtrees were chosen to be mutated to simulated the occurrence of a parallel mutation.

## Code availability

SCIΦN is available at https://github.com/cbg-ethz/SCIPhIN under a GNU General Public License v3.0 license.

## Data availability

The sequencing datasets were downloaded from the Sequence Read Archive with accession numbers SRA053195 and SRP044380.

## Competing interests

The authors declare no competing interests.

## Acknowledgements

The authors would like to thank Senbai Kang and Ewa Szczurek for discussions and correcting the base transitions in the sequencing error models. Part of this work was supported by SNSF Grant 310030_179518 (http://www.snf.ch).

## Author contributions

JK and JS developed the model, which was implemented by JS while the study was designed by JK, JS and NB. All authors drafted the manuscript and approved the final version.

## Supplementary Figures

**Supplementary Figure S1:**
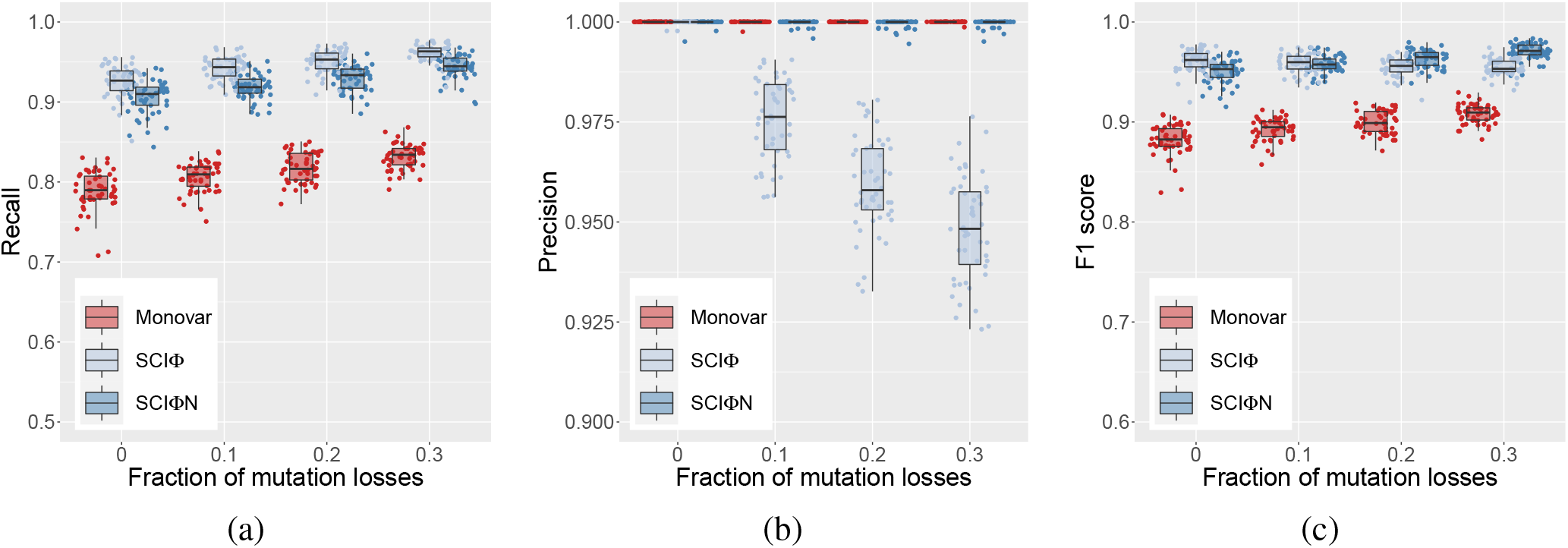
Effect of loss on single cell mutation calling.. The fraction of losses is increased (with no parallel mutations) to demonstrate their effect on the recall (a), precision (b) and the F1 score (c).

**Supplementary Figure S2:**
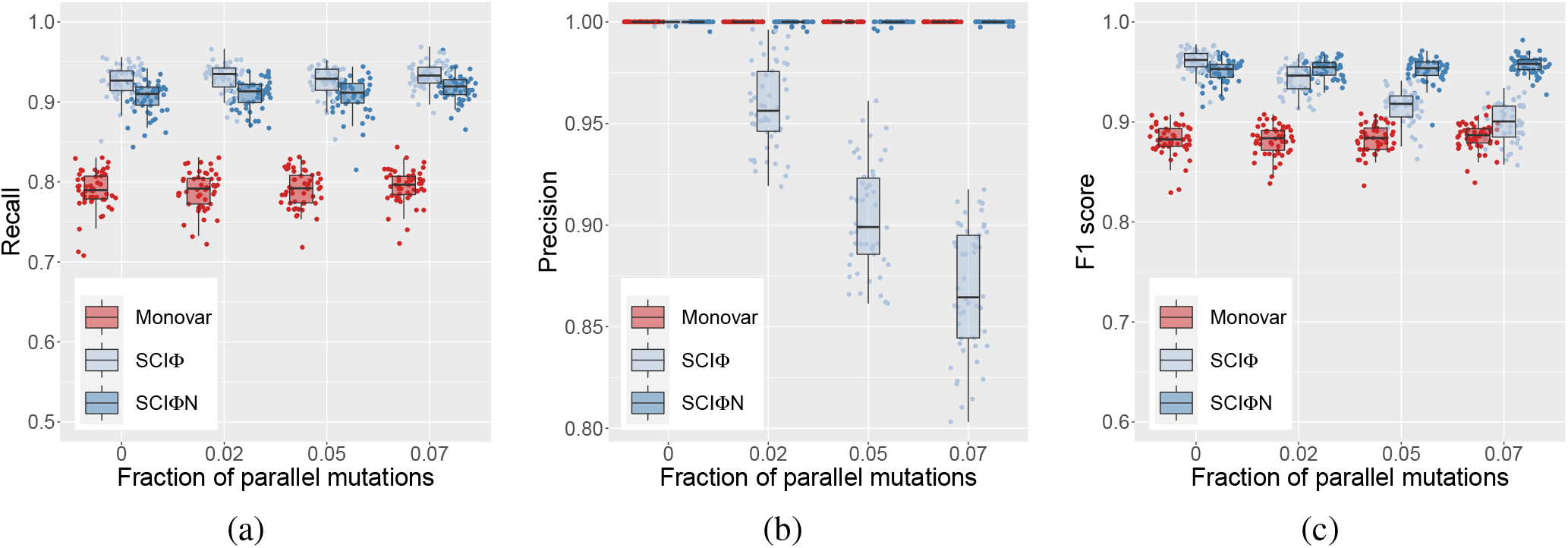
Effect of parallel mutations on single cell mutation calling. The fraction of parallel mutations is increased (with no losses) to demonstrate their effect on the recall (a), precision (b) and the F1 score (c).

**Supplementary Figure S3:**
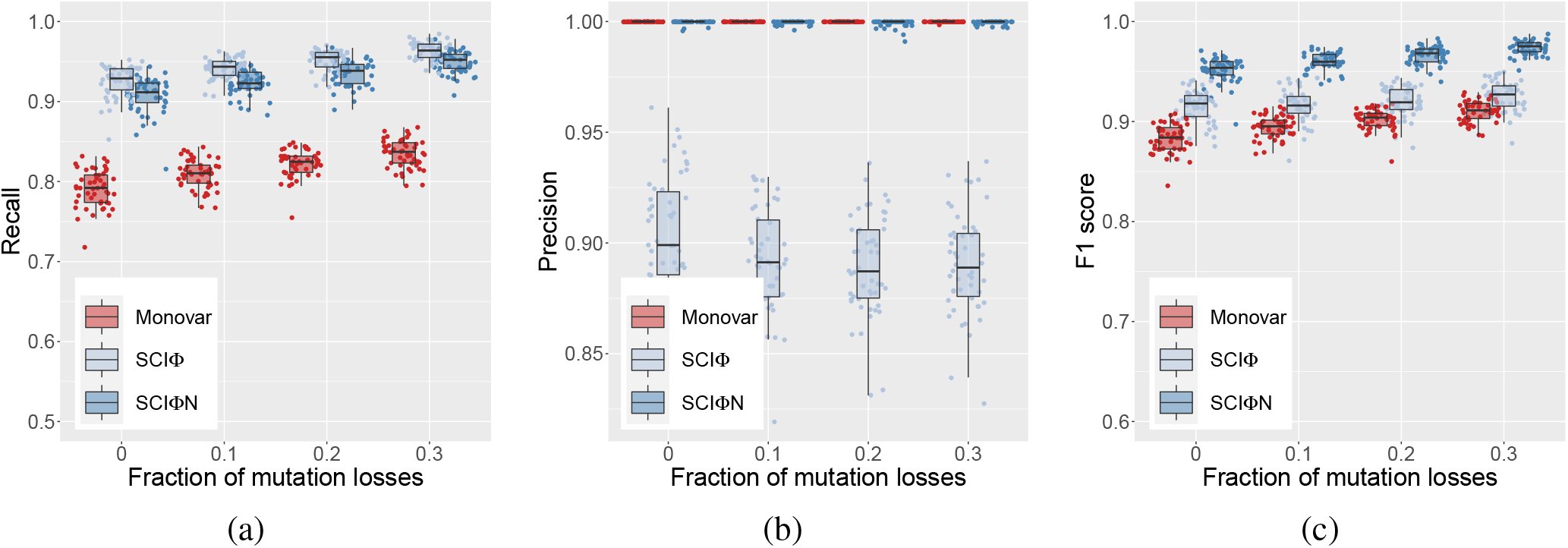
Effect of loss and parallel mutations on single cell mutation calling. The fraction of losses is increased (with 5% parallel mutations) to demonstrate their effect on the recall (a), precision (b) and the F1 score (c).

**Supplementary Figure S4:**
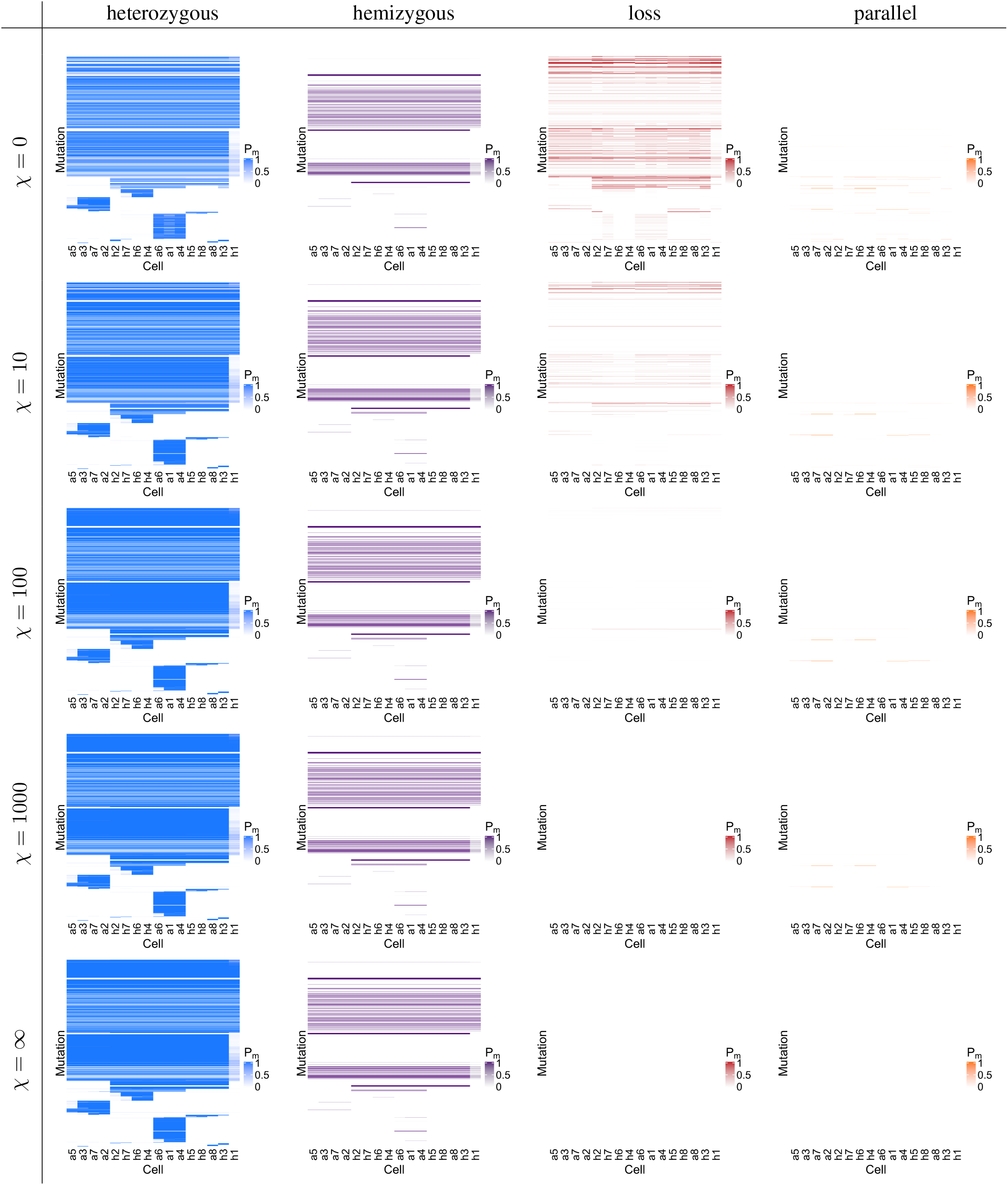
Mutation subtype calling on 16 breast cancer cells. The probability of mutation presence, *P_m_* displayed in Figure 3, broken down into components coming from heterozygous mutations, hemizygous mutations, loss of the wild type allele or loss of the mutation elsewhere in the tree, and from parallel mutations.

**Supplementary Figure S5:**
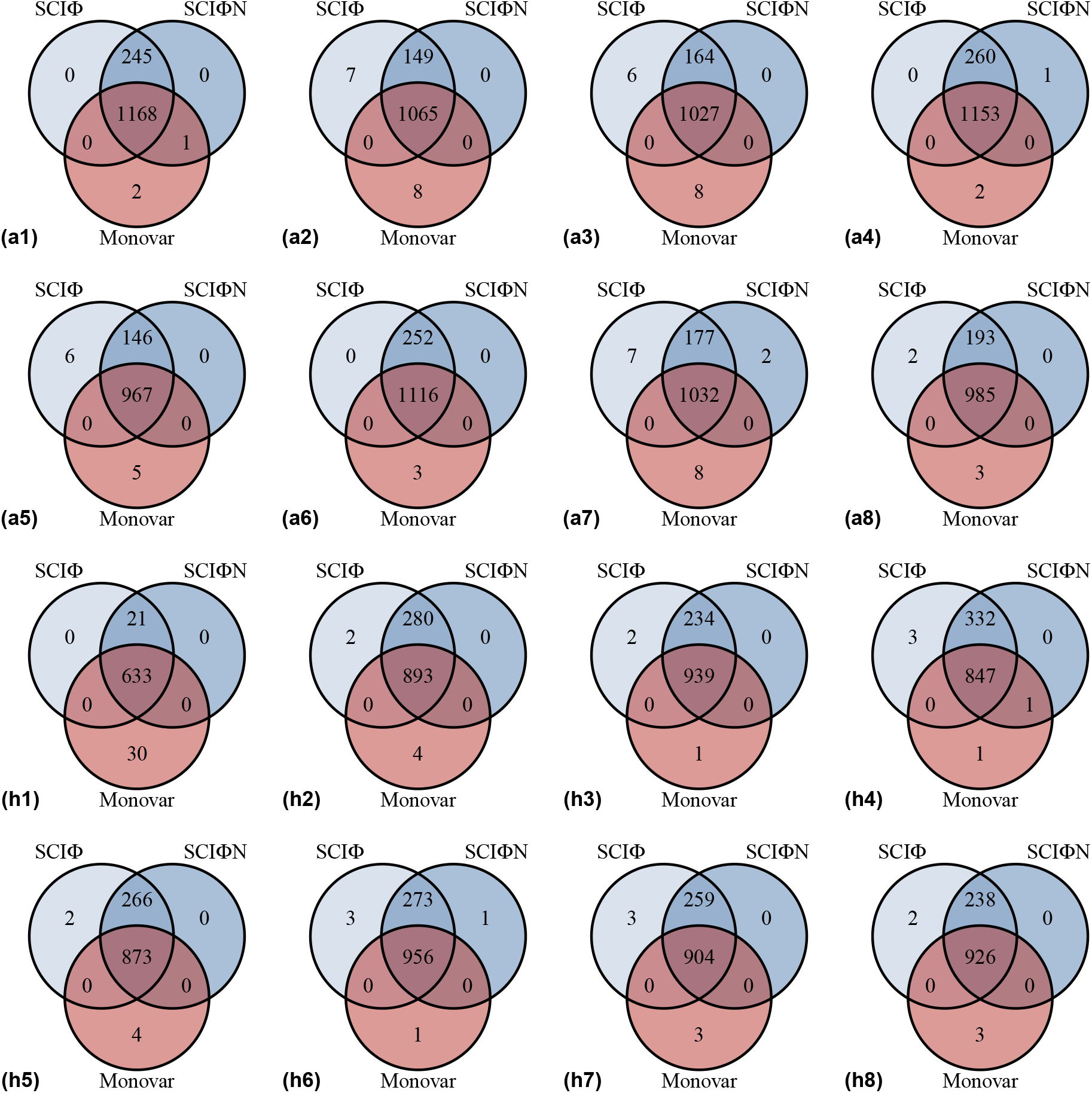
Concordance of mutation calls on 16 breast cancer cells. For each of the 16 single cells (labelled in brackets) we compare the called mutations from Monovar, SCIΦ and SCIΦN with penalisation *χ* = 100. Only loci identified by both Monovar and SCIΦN are included. For each cell, Monovar does not call mutations for loci with no coverage and these are therefore excluded from the comparison, even though SCIΦN can call mutations for such loci by sharing information from other cells through their phylogenetic relationships.

**Supplementary Figure S6:**
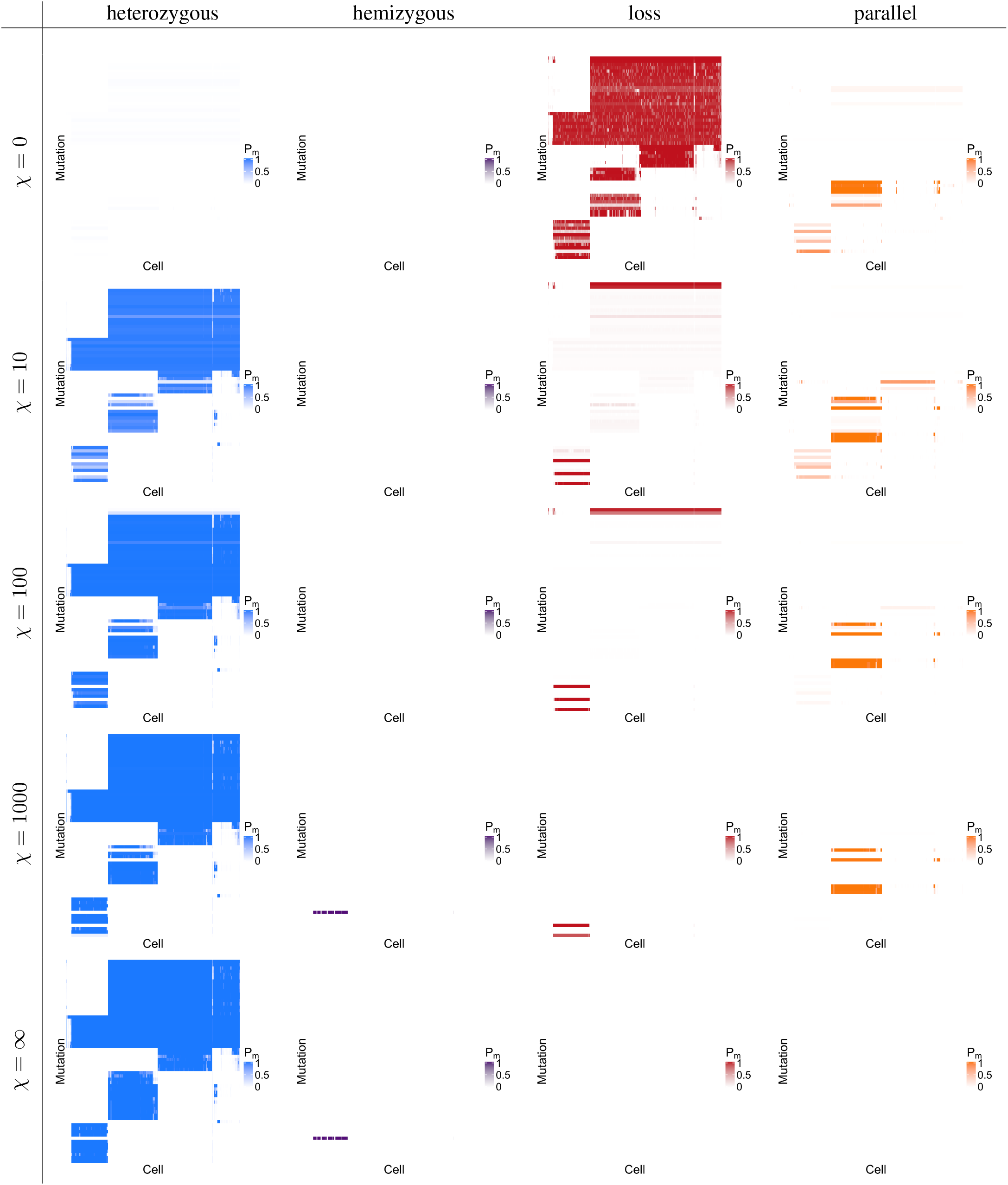
Mutation subtype calling on 255 acute myeloid leukemia cells. The break-downs of the probability of mutational presence from Figure 5, into components of heterozygous mutations, hemizygous mutations, lost mutations (where the wild type allele is lost or the mutation is lost elsewhere in the phylogeny) and parallel mutations.

## Footnotes

‡ N can abbreviate knight in chess/crosswords so the reading SCI-*finite* should indicate Single-Cell mutation Identification via *finite*-sites Phylogenetic Inference.

## Notes

### Competing Interest Statement

The authors have declared no competing interest.

